# Internal Protein Motions in Molecular Dynamics Simulations of Bragg and Diffuse X-ray Scattering

**DOI:** 10.1101/190496

**Authors:** Michael E. Wall

**Author notes:** Los Alamos National Laboratory technical release # LA-UR-17-27716.

## Abstract

**Synopsis:** A molecular dynamics simulation of diffuse X-ray scattering from staphylococcal nuclease crystals is greatly improved when the unit cell model is expanded to a 2×2×2 layout of eight unit cells. The dynamics are dominated by internal protein motions rather than rigid packing interactions.

**Abstract:** Molecular dynamics (MD) simulations of Bragg and diffuse X-ray scattering provide a means of obtaining experimentally validated models of protein conformational ensembles. This paper shows that, compared to a single periodic unit cell model, the accuracy in simulating diffuse scattering is increased when the crystal is modeled as a periodic supercell, consisting of a 2×2×2 layout of eight unit cells. The MD simulations capture the general dependence of correlations on the separation of atoms. There is substantial agreement between the simulated Bragg reflections and the crystal structure; there are local deviations, however, indicating both the limitation of using a single structure to model disordered regions of the protein and local deviations of the average structure away from the crystal structure. Although it was anticipated that a longer duration simulation might be required to achieve convergence of the diffuse scattering calculation using the supercell model, only a microsecond is required, the same as for the unit cell. Rigid protein motions only account for a small fraction of the variation in atom positions from the simulation. The results indicate that protein crystal dynamics can be dominated by internal motions rather than packing interactions, and that MD simulations can be combined with Bragg and diffuse X-ray scattering to model the protein conformational ensemble.

## Introduction

In X-ray diffraction from protein crystals, the sharp Bragg peaks are accompanied by diffuse scattering – streaks and cloudy features between the peaks. Diffuse scattering comes from imperfections in the crystal such as diverse protein conformations. Unlike the Bragg diffraction, which is only sensitive to the mean charge density, diffuse scattering is sensitive to the spatial correlations in charge density variations. Diffuse scattering therefore provides unique data for modeling protein conformational ensembles.

There is a longstanding interest in using diffuse scattering to validate MD simulations of protein crystals (1-7). Recent advances in computing now enable microsecond duration simulations of diffuse scattering (7) and Bragg diffraction (8, 9) that can overcome limitations seen using 10 ns or shorter MD trajectories (1, 4). In a microsecond simulation of a single staphylococcal nuclease unit cell (7), much of the agreement between the MD simulation and diffuse data is due to the isotropic component, a small-angle-scattering-like pattern seen for all protein crystals. Agreement with this component is significant as it consists of roughly equal contributions from solvent and protein (5, 7). The anisotropic component, which is about 10-fold weaker, agrees less well with the simulation (linear correlation of 0.35-0.43). This gap in accuracy between the isotropic and anisotropic components must be closed because the anisotropic component is richly structured and comes almost entirely from the protein, creating possibilities for validation of detailed models of protein motions. Accurate modeling of the anisotropic component is the key to unlocking the potential of diffuse scattering and MD simulations for modeling the conformational ensemble.

Wall et al. (7) noted that the simulation of a single unit cell might limit the accuracy of MD models of diffuse scattering, and suggested that simulating a larger section of the crystal with several unit cells might improve the accuracy. Here this idea is tested by constructing a periodic model of a 2x2x2 supercell of staphylococcal nuclease and performing a 5.1 microsecond duration MD simulation. The linear correlation of the anisotropic component of diffuse intensity computed from this simulation with the data is 0.68, indicating that the supercell simulation greatly increases the accuracy of the model. Analysis using Patterson methods suggests that distance dependence of the correlations is captured well. The mean structure factors from the simulation largely agree with the crystal structure; however, there are local deviations, suggesting a path to improve the MD model. The B factors from the simulation agree well with the crystal structure and improve on a TLS model. Similar to the unit cell simulation, the agreement of the supercell model with the data plateaus within a microsecond. This suggests that the simulation duration required to achieve convergence might become independent of the system size as it is increased beyond the length scale of the correlations. Finally, rigid body motions explain only a minority component of the dynamics, indicating that internal motions can be more important than packing dynamics in MD simulations of protein crystals.

## Methods

### Molecular dynamics simulation

A solvated crystalline model was created using PDB entry 1SNC. After stripping the waters, UCSF Chimera was used to add residues absent in the crystal structure. Using the context of the crystal structure as a guide, six missing residues at the N terminus were modeled as a beta strand, and eight missing residues at the C terminus were modeled as an alpha helix. A P1 unit cell of the protein and thymidine-3′-5′-bisphosphate (pdTp) ligand was built in UCSF Chimera using the P41 space group (4 copies per cell). The unit cell parameters were a=b=48.499 Å, c=63.430 Å, α=β=γ=90°. The system was extended to a 2x2x2 supercell in a 96.998 Å X 96.998 Å X 126.860 Å right rectangular box using Ambertools *PropPDB* (Fig. 1).

**Fig. 1.**
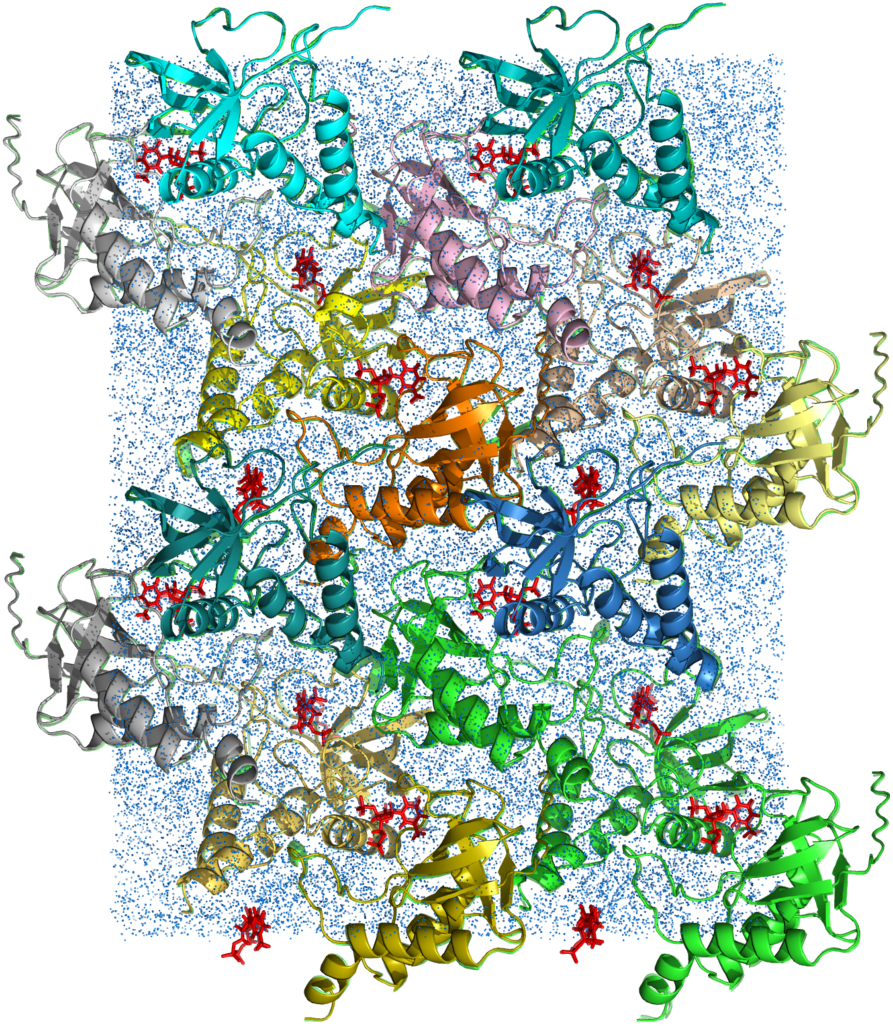
Staphylococcal nuclease supercell model. Eight unit cells containing four proteins each are arranged in a 2x2x2 layout. Protein chains are rendered as cartoons. The pdTP ligand is rendered as red sticks. Water atoms are indicated using speck-like blue spheres. The image was rendered using Pymol (https://pymol.org/).

Molecular dynamics simulations were performed using Gromacs (10) versions 5.0.2 (setup and first 4.1 μs) and 5.1.4 (extension to 5.1 μs). The protein topology was defined using *gmx grompp* with CHARMM 27 parameters (11, 12). The HIS protonation states from *grompp* were used without modification. 15,440 TIP3P water molecules were added using *gmx solvate*. To neutralize the system, 192 waters were replaced by Cl-ions, using *gmx genion*. CHARMM 27 compatible parameters for the pdTp ligand were obtained using the SwissParam server ((13), http://www.swissparam.ch/).

Simulations were performed using a constant NVT ensemble. NVT simulations are desired for crystalline simulations to enable comparisons of any calculated densities and structure factors to the crystal structure while avoiding difficulties introduced by drift of the unit cell parameters during the course of the simulation. The model after *gmx genion* showed large negative pressures when simulated via NVT. The standard approach for solvated systems of initially equilibrating the pressure using NPT simulations cannot be used, as this would change the box size and therefore the unit cell. The present approach is to iteratively perform energy minimization, NVT simulation, and solvation until obtaining a pressure near 1 bar. After several iterations, the number of water molecules was increased by 1,890 to 17,138. The mean pressure computed from the first 110 ns of the trajectory was 18 bar with a standard deviation of 130 bar, indicating that the procedure was successful.

The final system consisted of a total of 129,462 atoms. There were 32 copies of the protein, 32 copies of the pdTp ligand, 32 Ca^2+^ ions, 17138 water molecules, and 192 Cl-counterions.

For the production simulations, a time step of 2 fs was used, with LINCS holonomic constraints on all bonds. Neighbor searching was performed every 10 steps. The PME algorithm was used for electrostatic interactions, with a cutoff of 1.4 nm. A reciprocal grid of 64 x 64 x 80 cells was used with 4th order B-spline interpolation. A single cut-off of 1.4 nm was used for Van der Waals interactions. Temperature coupling was done with the V-rescale algorithm. The protein-ligand complex was treated as a separate temperature group from the rest of the atoms. Periodic boundary conditions were used. Trajectory snapshots were obtained every 2 ps in Gromacs .xtc format. The first 110 ns of the trajectory was dedicated to initial setup and equilibration. An initial production run then extended the duration to 1.1 μs. Subsequent extensions were performed to 5.1 μs, in 1 μs increments. The size of the equilibration trajectory is 27 GB, and the rest are each 245 GB. Each microsecond took about 2-4 weeks to complete on LANL Institutional Computing machines, depending on availability of cycles.

### Simulated diffuse intensity

The diffuse intensity was calculated for 100 ns sections of the MD trajectory. Each section was divided into 200 chunks, which were processed in parallel across 10 nodes of an Intel Xeon E5-2660_v3 cluster. Prior to performing the calculation, each snapshot of the trajectory was aligned to the crystal structure using the gromacs .tpr structure file. To do this, the .tpr file was converted to a .pdb file using gmx editconf. The .pdb file was processed to ensure the coordinates reflected the connectivity of the molecules (*gmx trjconv –pbc mol*). The alignment was performed using the processed .pdb file as the reference structure (*gmx trjconv -fit translation –pbc nojump*).

Each chunk of sampled structures was processed using the previously described Python script *get_diffuse_from_md.py* (7) to calculate the diffuse intensity to 1.6 Å resolution. The calculation of the diffuse intensity *D*_*md*_(*hkl*) uses Guinier’s equation (14):

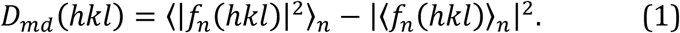

In the script, the structure factor, *f*_*n*_(*hkl*), for each sample *n* is calculated at Miller indices *hkl* using the iotbx package in the computational crystallography toolbox (CCTBX) (15). The script was modified to accept input of an externally supplied unit cell specification using the PDB CRYST1 format. Specifying the P1 unit cell from the crystal structure yields the diffuse intensity *D*_*md*, 1×_(*hkl*) sampled on the Bragg lattice, only at integer *hkl* values. Specifying the P1 supercell dimensions in the CRYST1 record yields *D*_*md*, 2×_(*hkl*), which is sampled twice as finely, at *hkl* values that are multiples of ½. Averages for longer sections of the trajectory were accumulated from averages of the smaller chunks.

To decompose the diffuse intensity into isotropic and anisotropic components, reciprocal space was subdivided into concentric spherical shells, each with a thickness equal to the voxel diagonal. The discretely sampled isotropic intensity *D*_*md*_(*s*_*n*_) was calculated as the mean intensity at scattering vector *s*_*n*_ at the midpoint of each shell *n*. The anisotropic intensity 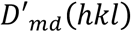 was then calculated at each lattice point *hkl* by subtracting the isotropic intensity *D*_*md*_(*s*_*hkl*_) from the original signal *D*_*md*_(*hkl*). The value of *D*_*md*_(*s*_*hkl*_) at scattering vector *s*_*hkl*_ in the range (*s*_*n*_, *s*_*n*+1_) was obtained by cubic B-spline interpolation of *D*_*md*_(*s*_*n*_) (previous anisotropic intensity calculations made use of linear interpolation (7); results using either interpolation method were similar in the present case, though the spline is generally preferred for increased accuracy). The same method was used to obtain isotropic (*D*_*o*_(s_n_)) and anisotropic 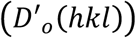 components of the experimentally observed diffuse intensity.

Because the experimental diffuse intensity shows symmetry consistent with the P4_1_ symmetry of the unit cell, the P4/*m* Patterson symmetry (corresponding to the P4_1_ unit cell symmetry) was enforced by replacing each *D*_*md*_(*hkl*) value with the average over all symmetry equivalent *hkl* positions in the map.

### Simulated structure factors and average structure

Averages of *A*_*n*_(*hkl*) were computed along with the diffuse intensity. To obtain structure factors for comparison with the crystal structure, the P41 unit cell CRYST1 record was used in lieu of the P1 unit cell or supercell record. The real space correlation coefficient (RSCC) was evaluated using the MolProbity validation tool in Phenix (16), using PDB entry 4WOR and the intensities *I*_*md*_(*hkl*), computed as the square of the mean *f*_*n*_(*hkl*). The errors in the intensities were calculated as the square root of the intensities. Prior to calculating the RSCC, a molecular replacement search using CCP4 *molrep* was used to determine the placement of the protein in the unit cell. The average structure from the simulation was computed by using *phenix.refine* to refine the crystal structure against the *I*_*md*_(*hkl*). The *phenix.refine* option *apply_overall_isotropic_scale_to_adp=false* was used to remove the bulk-solvent scaling component from the B factors. For comparison, the B factors from a TLS model were obtained by refinement against the experimental Bragg data using the *refine.adp.tls="chain A" strategy=tls* in *phenix.refine*.

### Image processing and diffuse data integration

Experimental diffuse scattering data from Wall *et al*. (17) were used for validation of the simulations. These data were collected on a custom CCD detector configured in an anti-blooming mode in which charge was drained away from overflow pixels (18). The data were processed using Lunus software for diffuse scattering ((19), https://github.com/mewall/lunus). Indexing was performed using image numbers 1, 20, and 40 from the rotation series. Rather than limiting the observations *D*_*o*_(*hkl*) to integer values of the Miller indices *hkl*, as was done for the single unit cell MD simulations (7), data were sampled twice as finely, both at Miller indices and at the mid-points between. The two-fold sampling yields data that correspond precisely to the reciprocal lattice of the 2x2x2 supercell from the MD model. To place the model and data in an equivalent orientation, the data were reindexed by applying a 180 degree rotation about the *h*-axis. Prior to integration, images were mode filtered to reject the Bragg peak signal (18, 19). The kernel for the mode filter was a 15 pixel x 15 pixel square, with frequency statistics evaluated in 1 ADU bins.

The accuracy of Lunus was improved by shifting from integer to floating point arithmetic for the polarization correction and sold angle normalization, which were combined into a single step. The increase in accuracy is small for strong diffraction images such as those used here but is substantial for weaker diffraction images, e.g., including pixels with fewer than 10-100 photon counts. In addition, as mentioned above (Diffuse scattering computation), a cubic B-spline method was implemented in Lunus to improve calculation of the anisotropic intensity.

A collection of helper scripts was added to Lunus to enable diffraction images to be processed and integrated in parallel. The helper scripts produce shell scripts with Lunus workflows to perform image processing, integration, and merging of the data, and a CCTBX (15) workflow to index the data and obtain a transformation to map pixels in diffraction images to fractional *hkl* values in reciprocal space. The scripts were executed in parallel on 12 nodes of a 32-core Intel Haswell cluster; the 96 1024x1024 images of diffraction from staphylococcal nuclease could be processed in one minute. Real-time parallel processing of diffuse scattering from single-crystal synchrotron datasets is therefore now possible using Lunus.

All Lunus revisions, including helper scripts for parallel processing, have been committed to the github repository, https://github.com/mewall/lunus.

As for the calculated *D*_*md*_(*hkl*) the P4/*m* Patterson symmetry was enforced for the experimental *D*_*o*_(*hkl*). The isotropic and anisotropic components *D*_*o*_(*s*_*n*_) and 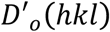 were calculated from *D*_*o*_(*hkl*), like for *D*_*md*_(*s*_*n*_) and 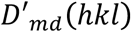 (Diffuse scattering computation). The correlation coefficient *r*_*oc*_ was used to compare the total calculated diffuse scattering, *D*_*md*_(*hkl*), to the experimental data, *D*_*o*_(*hkl*), and used the correlation coefficient 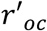 to compare the anisotropic component, 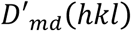 to the experimental data, 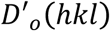.

### Simulated diffraction images

Diffraction images were simulated using methods similar to the data integration, except instead of the datasets being compiled from the pixel values, pixel values were obtained from 3D datasets. An indexing solution was obtained as in the data integration, and a template image was provided to determine the crystal orientation. Each pixel was mapped to a fractional Miller index, and the pixel value was calculated as a sum of intensities at the eight nearest grid points in the dataset, in proportion to the distance to the center of the grid point along each axis in the space of Miller indices. The method was implemented in a Lunus python script, *simulate_diffraction_image.py* using CCTBX methods.

### Patterson maps

Diffuse Patterson maps were created by Fourier transforming diffuse intensities. Symmetrized anisotropic diffuse intensities were output in *hklI* text format using Lunus *lat2hkl*, and were converted to .mtz format using *phenix.reflection_file_converter*. Fourier transforms were computed using the PATTERSON *fft* method (20) in the CCP4 suite (21). A Patterson map of the anisotropic component of the Bragg reflections was computed, for comparison to the diffuse Patterson maps. The anisotropic component of Bragg reflections was calculated using Lunus *anisolt*, after conversion of the reflections from .mtz format to hklI text using CCP4 *mtz2various*. The intensities were than converted back to .mtz using *phenix.reflection_file_converter*, and the Patterson obtained using *fft*, as mentioned above.

### Rigid-body rotation analysis

Snapshots of alpha carbon positions from the supercell were obtained every 40 ps and were translationally aligned to the structure in the .tpr file used for the 110-1100 ns simulation. Rotation matrix analysis was performed using 32 runs of *gmx rotmat*, using the snapshots for each of the 32 copies of the protein as inputs. The *gmx_rotmat.c* source code was edited to add outputs of the root mean squared deviation (RMSD) of coordinates between the snapshot and the reference structure both before and after alignment, in addition to the usual output, the elements of the rotation matrix. A custom python script was written to compute standard deviations of Euler angles from the .xvg rotation matrix element output of *rotmat*, and to accumulate trajectory-wide RMSDs by calculating the square root of the average squared RMSD for individual snapshots.

## Results

MD simulations were performed using a solvated supercell model of crystalline staphylococcal nuclease consisting of eight unit cells in a 2x2x2 layout (Fig. 1, Methods). The total simulation duration was 5.1 μs. Trajectories were obtained for the following segments, sampled every 2 ps: the initial equilibration, 0-110 ns; 110– 1100 ns; 1100-2100 ns; 2100-3100 ns; 3100-4100 ns; and 4100-5100 ns.

Diffuse intensities were calculated from the trajectories in 100 ns sections, and the agreement with the data was assessed using the anisotropic component (Methods). In the 100 ns immediately following the equilibration, the linear correlation between the simulation and the data is 0.62. Correlations range between 0.59 and 0.62 for subsequent sections, through 1100 ns (Fig. 2, dashed boxes).

**Fig. 2.**
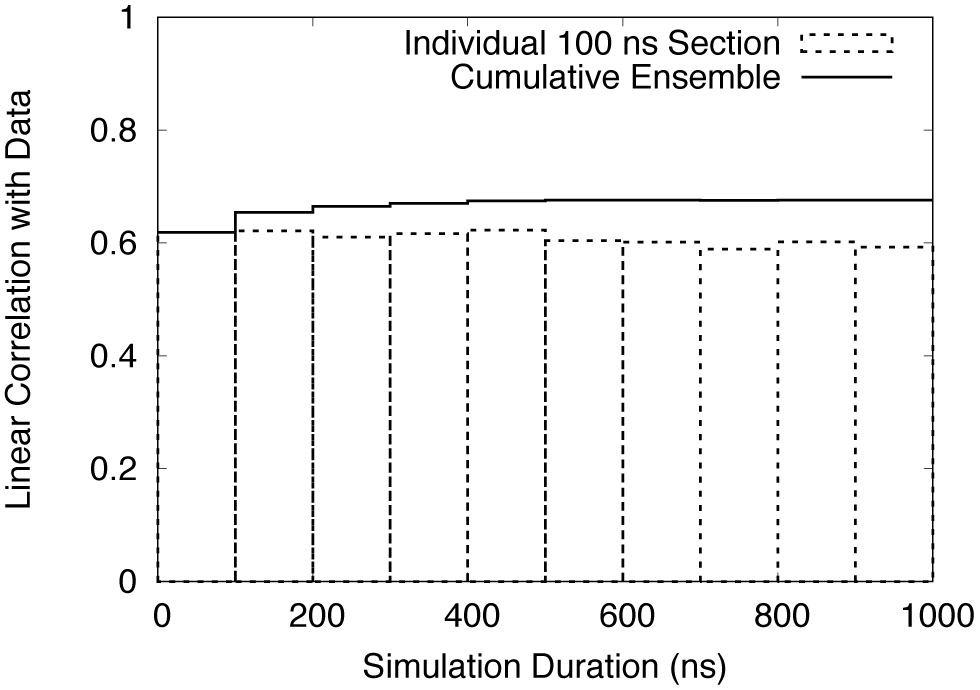
Linear correlation between the simulated and experimental diffuse intensity, evaluated for subsequent 100 ns sections of the MD trajectory (dashed boxes) and cumulatively (solid line). The cumulative plot plateaus at a value of 0.68.

The correlation of the running average of the diffuse intensity with the data increases steadily from 0.62 to 0.68 from 110-700 ns; it remains at 0.68 through 1100 ns (Fig. 2, solid line). Beyond 1100 ns, the agreement decreases: the correlation for 100 ns sections between 1100-5100 ns ranges from 0.56 to 0.60; the correlations for the accumulated 1100-2100 ns, 2100-3100 ns, and 3100-4100 segments are all 0.65; and the correlation for the 4100 ns-5100 ns segment is 0.63.

Simulated and experimentally derived diffraction images look similar (Fig. 3). There is good correspondence between the shapes of cloudy features in the simulation and the experimental data at all but the lowest resolutions. There are some large differences in the strengths of the features; for example, the large, intense red feature in the bottom half of the simulation image (Fig. 3A) is weaker than the corresponding feature in the experimentally derived image (Fig. 3B).

**Fig. 3.**
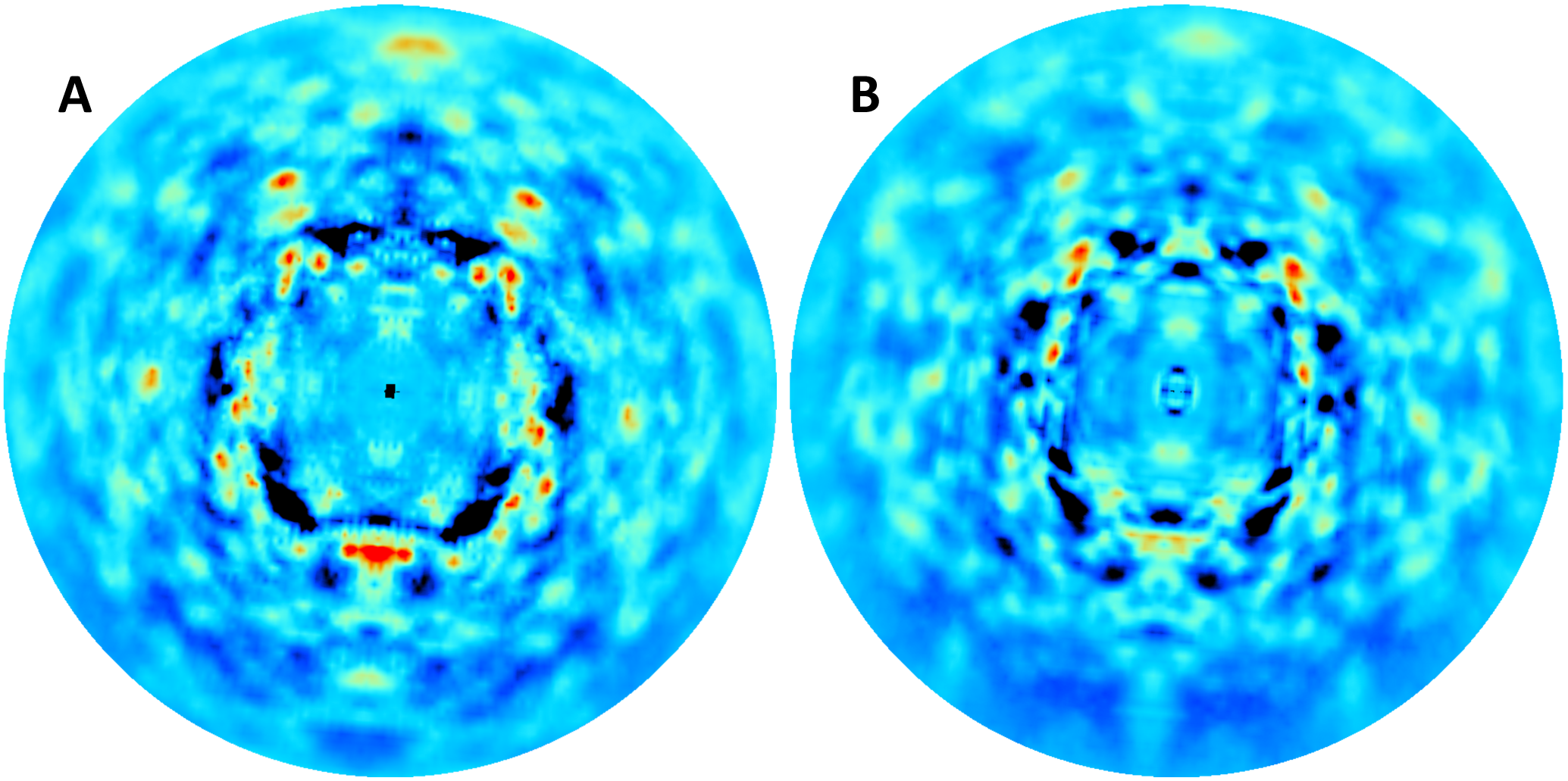
Comparison of simulated diffraction images calculated from the simulated (Left panels) and experimental (Right panels) three-dimensional diffuse intensities. The crystal orientation corresponds to the first diffraction image in the rotation series. The display is truncated at 1.6 Angstroms. The mean pixel value at each scattering vector was subtracted, and then the minimum value in the image was subtracted, prior to visualization. The images were displayed using the rainbow colormap in Adxv (38), with a pixel value range arbitrarily chosen to highlight the similarities.

The correlation between the simulation and the data is substantial over a wide resolution range (Fig. 4, solid steps), consistent with the range over which the diffuse features look similar in Fig. 3. Above 10 Å resolution, the correlation is at least 0.52 in all of the 32 resolution shells, and is above 0.6 in all but three. Below 3 Å resolution, the agreement between the simulation and data is substantially less than the agreement between symmetrized and unsymmetrized datasets, indicating there is room for improvement in modeling the data in this resolution range. Above 2 Å resolution, the agreement between the data and simulation is higher than the degree to which the symmetry is obeyed, suggesting that the symmetry averaging might have eliminated some systematic error in the data. Below 10 Å resolution, the agreement becomes very small (with the exception of a low resolution outlier), and the CCsym values also become very small, suggesting that the data were not accurately measured in the neighborhood of the beam stop.

**Fig. 4.**
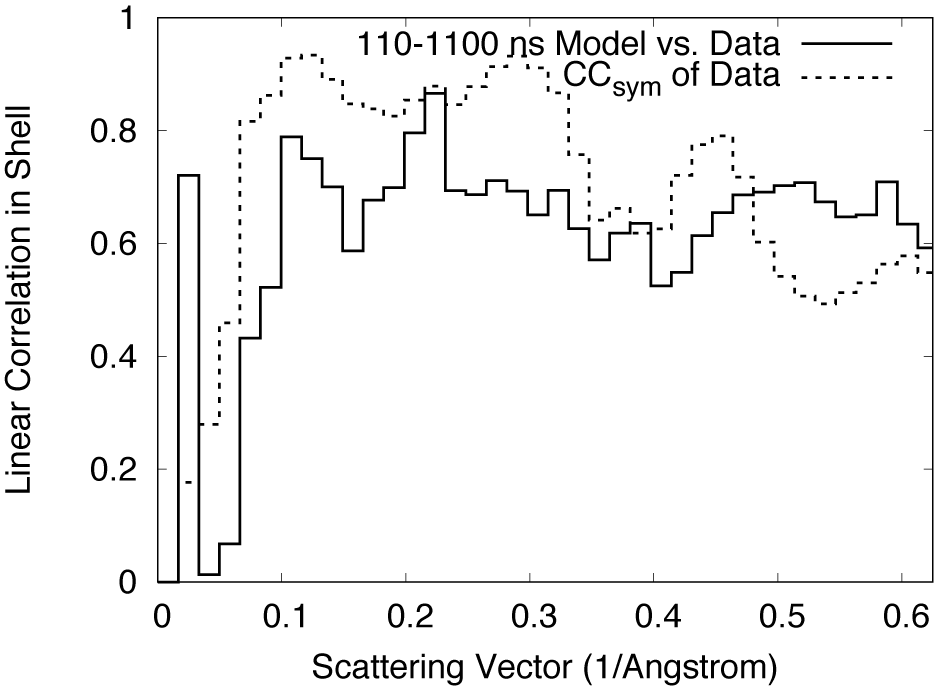
Resolution-dependent agreement between simulation and data (solid line), compared to the self-consistency of the data, as assessed using the expected P4/*m* symmetry (dotted line).

Real-space comparisons of the Patterson function of the charge density variations (Methods) show that both the simulation and the data exhibit similar modulation with distance (Fig. 5). The amplitude of the variations is especially similar in the *X*=0 section (Figs. 5A,B). In the *Z*=0 section, the simulation has higher amplitude variations than the data at longer distances (Figs. 5D, E), indicating that the correlations within this plane are stronger in the simulation than in the data. The linear correlation between the Pattersons computed from the simulation and the data is 0.70. The Patterson computed from the Bragg data (Figs. 5C, F) shows much higher amplitude features at long distances, indicating a longer length scale of correlations for the mean than for the variations in charge density.

**Fig. 5.**
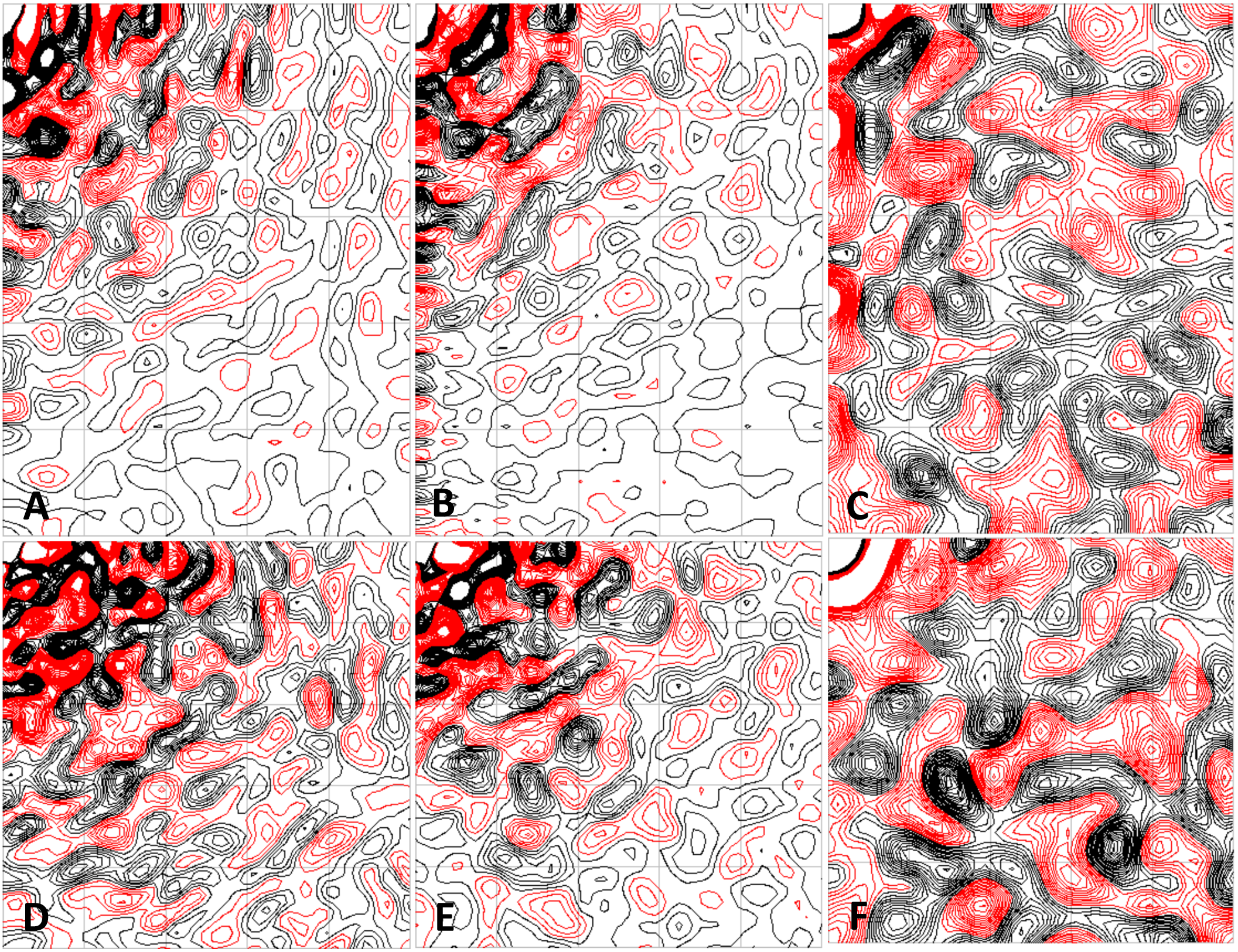
Comparison of the Fourier transform of the simulated diffuse intensity (left panels) with that of the experimental data (middle panels) and the Bragg data (right panels). Only the anisotropic component is used to calculate the transforms. Positive contours are in black, negative in red. Contours are at every 0.5-sigma between 0 and 10-sigma in the diffuse data and are adjusted to be equivalent with respect to sigma/Imax in other panels. (A) *X*=0 section, simulation; (B) *X*=0 section, diffuse data; (C) *X*=0 section, Bragg data; (D) *Z*=0 section, simulation; (E) *Z*=0 section, diffuse data; (F) *Z*=0 section, Bragg data. The plots were produced using CCP4 mapslicer (21).

The real-space correlation coefficient (RSCC) computed using the crystal structure and the calculated Bragg reflection amplitudes from the simulation (Methods) is high in most regions (Fig. 6, purple line); however, there are especially large dips (<0.6) for residues 6-8 at the N terminus, and for residues 46-52. The average RSCC for all residues is 0.80. Regions of the crystal structure with high B factors (Fig. 6, blue line) include both the N terminus and a previously observed disordered loop in the crystal structure at residues 44-50 (22). The overall RMSD of atom positions between the simulation average structure (Methods) and the crystal structure is 0.7 Angstroms, with high deviations concentrated in local regions of the protein (Fig. 6, yellow line). The residue-wise composite B factors from the crystal structure and the simulation average structure are very similar (Fig. 7A,B); the linear correlation between the two is 0.94. By comparison, the residue-wise composite B factors from a TLS model of the crystal structure underestimate the disorder in the N terminus and in the flexible loop (Fig. 7C); the linear correlation between the B factors derived from the individual ADP vs TLS model is 0.89.

**Fig. 6.**
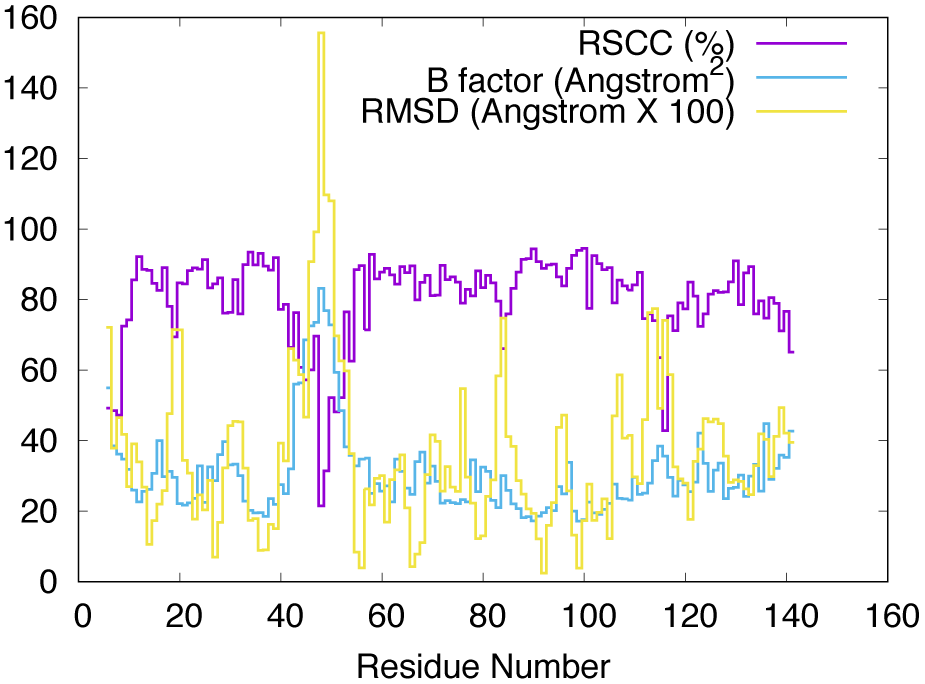
Residue-wise comparisons between the simulated structure factors and the crystal structure. The real space correlation coefficient (purple line) is computed using the crystal structure and the simulated Bragg reflections. The isotropic B factors (blue line) are taken from the crystal structure. The RMSD of atomic coordinates (yellow line) is computed between the average structure from the simulation and the crystal structure.

**Fig. 7.**
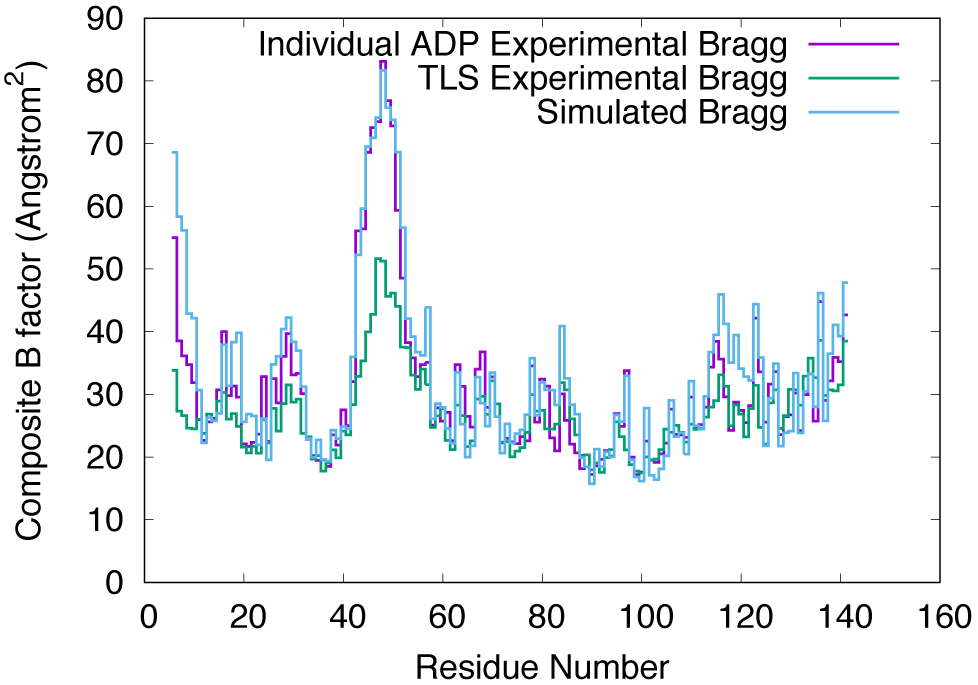
Residue-wise comparison between the B factors from the crystal structure (purple), MD simulated average structure (cyan), and a TLS model of the crystal structure (green).

## Discussion

The 0.68 correlation with the anisotropic component of the diffuse data is much higher than has been previously achieved using MD simulations. The present simulation used a supercell model whereas previous simulations used a unit cell model of the crystalline protein. Supercell models are expected to be more accurate, as they more realistically describe interactions across unit cell boundaries. Larger supercell models might further increase the accuracy, and can yield detailed models of large-scale motions that have been studied using simpler models of protein diffuse scattering, such as acoustic crystal vibrations in ribosome crystals (23), coupled rigid-body motions in lysozyme crystals (24), and liquid-like motions with long correlation lengths in lysozyme (25) and calmodulin crystals (26).

In addition, compared to the previous simulation, the present simulation included residues theoretically modeled at the N and C terminus. To determine the role of including the extra residues in achieving the increased correlation, a MD simulation of a single unit cell was performed using the extended model (unpublished). The correlation between the simulated and experimental diffuse intensity within the first microsecond was 0.42 compared to the previous correlation of 0.35-0.43 (7). The supercell model therefore accounts for the increased accuracy of the simulated diffuse scattering.

Like in the supercell simulation, the residue-wise B factors of the average structure from the single unit cell simulation are similar to the crystal structure (Supporting Fig. S1A); the linear correlation between the two is 0.95, compared to a correlation of 0.94 for the supercell simulation. A similar high agreement with crystallographic B factors was seen in MD simulations of a 3x2x2 supercell P1 hen egg-white lysozyme crystals (9). The RSCC values computed between the crystal structure and the unit cell simulation also are similar to those for the supercell simulation (Supporting Fig. S1B); the average RSCC for all residues is 0.82 for the unit cell simulation, compared to 0.80 for the supercell simulation. Compared to the unit cell simulation, the supercell simulation therefore specifically improves the model of structure variations, but not the model of the average structure.

The maximum agreement with the data was achieved within the first 1100 ns of the simulation. The rate of convergence to the maximum is similar to what was seen for the previously published unit cell simulation (7). Because the diffuse intensity is only sensitive to the two-point correlations in the variations, this result suggests that the most important motions in the present MD simulation for explaining the experimental data are correlated on a shorter length scale than the unit cell. The short length scale of correlations is supported by the comparison between the Patterson computed from the Bragg and diffuse data, which reveal that the diffuse Patterson is attenuated at long distances (Fig. 5). The simulation duration required to achieve a similar accuracy is therefore expected to be nearly independent of the system size. Prior to this study, the expectation was that supercell simulations would require a longer duration than unit cell simulations to converge, as larger systems involve motions on longer length scales, which are generally slower. It is still possible that more accurate MD models of larger scale motions would yield an even higher correlation with the data, at the cost of longer simulation times. However, the finding that reasonably accurate MD models of larger systems do not require longer simulation durations is surprising.

There are strong similarities between the Patterson maps calculated from the simulated and experimental diffuse intensities (Fig. 5). The overall modulation of the diffuse Patterson is especially similar between the simulation and the data, indicating that the distance dependence of the correlations is captured well by the MD simulation. The attenuation of the Patterson along the *a* and *b* lattice vectors is more pronounced in the data than in the simulation, however, indicating a longer length scale of correlations in the simulation than in the data within the *Z* = 0 plane (Fig. 5C,D).

If the simulation perfectly described the experimental system, the agreement would be expected either to increase or to plateau at long times, depending on how accurately the data were measured. The agreement beyond 1100 ns decreased, however, indicating a divergence of the simulation away from the data. While it is possible that the decrease is transient and that running the present simulation for longer would eventually lead to an increase in the agreement, the simplest explanation is that the MD model is deviating from the experimental behavior at long times.

Dips in the residue-wise RSCC plot (Fig. 6, purple line) indicate regions where the simulated charge density locally deviates from the crystal structure. Dips at the N terminus and at residues 46-52 correspond to high B factor regions (Fig. 6, blue line), and might reflect the intrinsic difficulty of capturing discrete conformational variability using B factors (27). Dips in the RSCC also correspond to regions of high RMSD between the simulation average structure and the crystal structure (Fig. 6, yellow line). These include not only the high B factor regions, but also many regions with lower B factors. The low B factor regions with high RMSDs indicate where the atom positions from the simulation locally deviate from the crystal structure. The discrepancies in these regions would be especially good targets for improving the MD model.

There are a number of specific routes to improving the MD model. Missing residues at the N and C terminus could be modeled more accurately, using more of the context from the crystal structure. The 2x2x2 supercell could be extended to an even larger supercell. Additional compounds found in the mother liquor could be added to the model (e.g. 23% 2-methyl-2,4-pentanediol (MPD)), and the ionic strength of the solvent could be more accurately modeled; the current model only includes water and neutralizing counter-ions. There might be inaccuracies in the MD force fields; importantly, the accuracy of the MD models should now be high enough to enable improvement of force fields using crystallography data, just as validation using NMR data (28) has led to improvement in MD force fields (29-31). Time-averaged ensemble refinement produces models that are closer to the crystal structure (32) and might be used in combination with diffuse data to generate more accurate conformational ensembles.

Recent solid state NMR (ssNMR) experiments combined with crystalline protein simulations (33-35) create opportunities for joint validation of MD simulations using crystallography and NMR. A ssNMR + MD study of protein GB1 (35) showed reasonable agreement between 200 ns MD simulations and the data for longitudinal relaxation rates and chemical shifts, with lower agreement for the transverse relaxation rates. A ssNMR + MD study of ubiquitin (33, 34) attributed the transverse relaxation rates to rigid body rotations of whole proteins in the crystal lattice, with amplitudes of 3-5 degrees extracted from the simulations. To assess the importance of rigid body rotations in the present staphylococcal nuclease simulation, snapshots of each of the 32 copies of the protein were rotationally aligned to a reference structure (Methods). Standard deviations of Euler angles were mostly in the 1-2 degree range, with individual values as low as 0.8 degrees and as high as 2.3 degrees (Fig. 8A). The RMSD of coordinates between the snapshots and the reference structure decreased after the alignment, but only by 10-20% for most copies of the protein, with a minimum of 8% and a maximum of 28% (Fig. 8B). Therefore rigid body rotations are not a substantial component of the dynamics in the present simulations.

**Fig. 8.**
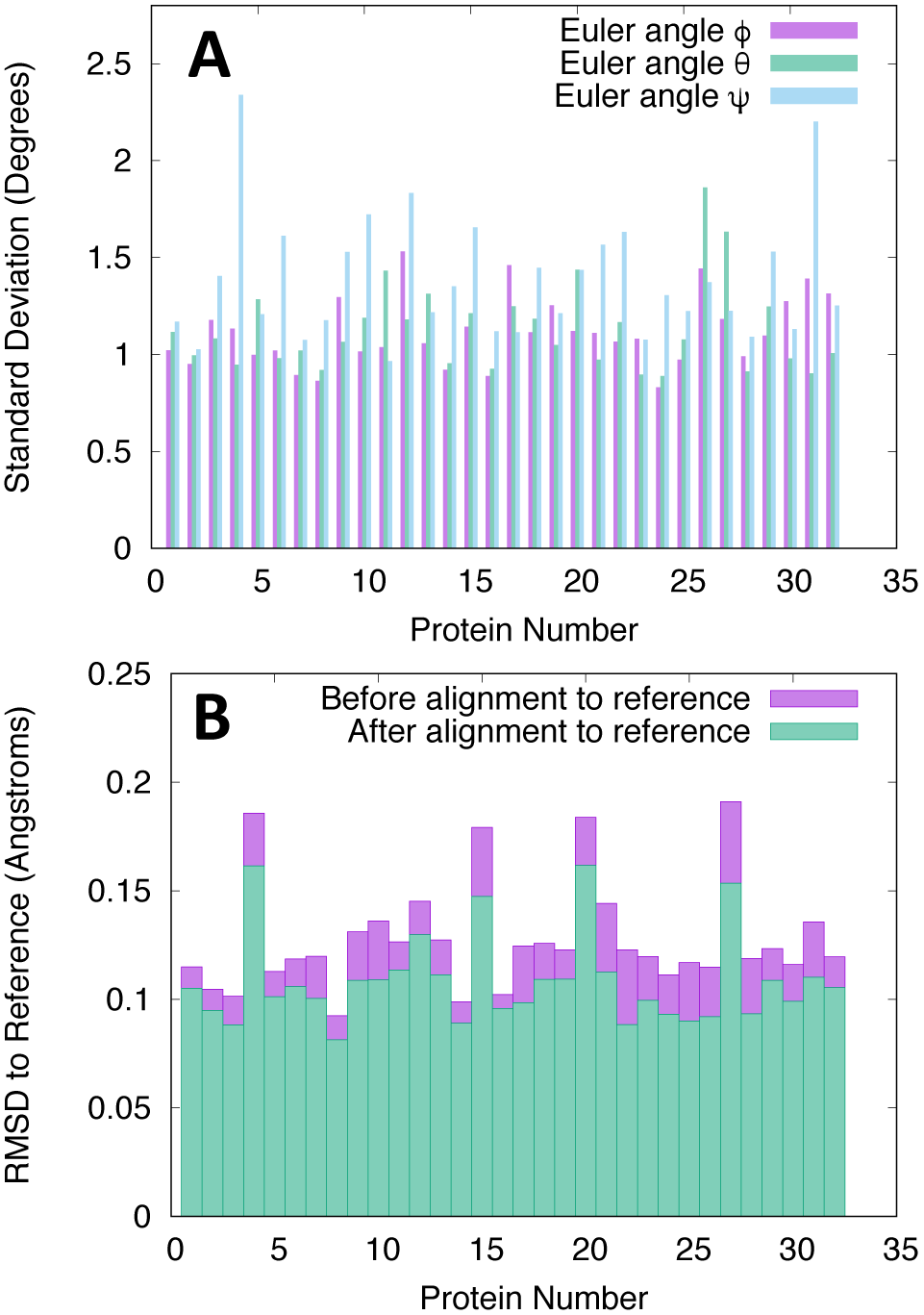
Analysis of the MD trajectory in terms of rigid-body motions of whole proteins. (A) Standard deviations of Euler angles that optimally align protein snapshots to the reference structure. (B) RMSDs of coordinates computed before and after aligning protein snapshots to the reference structure.

Further investigation of the rotational matrix fit for protein numbers 4 and 31, which have ψ angle SDs of 2.3 and 2.2 degrees, respectively, revealed a pitfall in rotational analysis. Visual inspection of the trajectories for these protein numbers revealed a conformational change in the flexible loop around residues 42-54 during the first microsecond (Fig. 9). When the tip of the loop was removed (residues 46-52, rendered using sticks in Fig. 9), the fitted value of ψ decreased by 0.7 degrees for each of the protein copies. This means that the rotational matrix fit does not solely report on rigid body motions, as is commonly assumed; therefore caution is warranted when using rigid-body motions models to interpret MD simulations. It would be interesting to perform a similar analysis of crystalline ubiquitin MD trajectories (33, 34), to see whether the variations in rotational fits correspond to rigid body rotations, as assumed, or whether they instead reflect internal motions. Overall the analysis here, including the improved modeling of the B factors using the MD model compared to a TLS model (Fig. 7), highlights the importance of internal motions and suggests a more minor role for independent rigid body translations (36) or rotations (37) in protein diffuse scattering. It will be important to determine whether rigid body motions are important for other proteins, especially proteins that are stiffer than staphylococcal nuclease. Studies that combine crystallography, ssNMR, and MD simulations to develop accurate models of crystalline protein dynamics with multiple points of validation are strongly motivated, to reveal the mechanisms of variation that really occur in protein crystals.

**Fig. 9.**
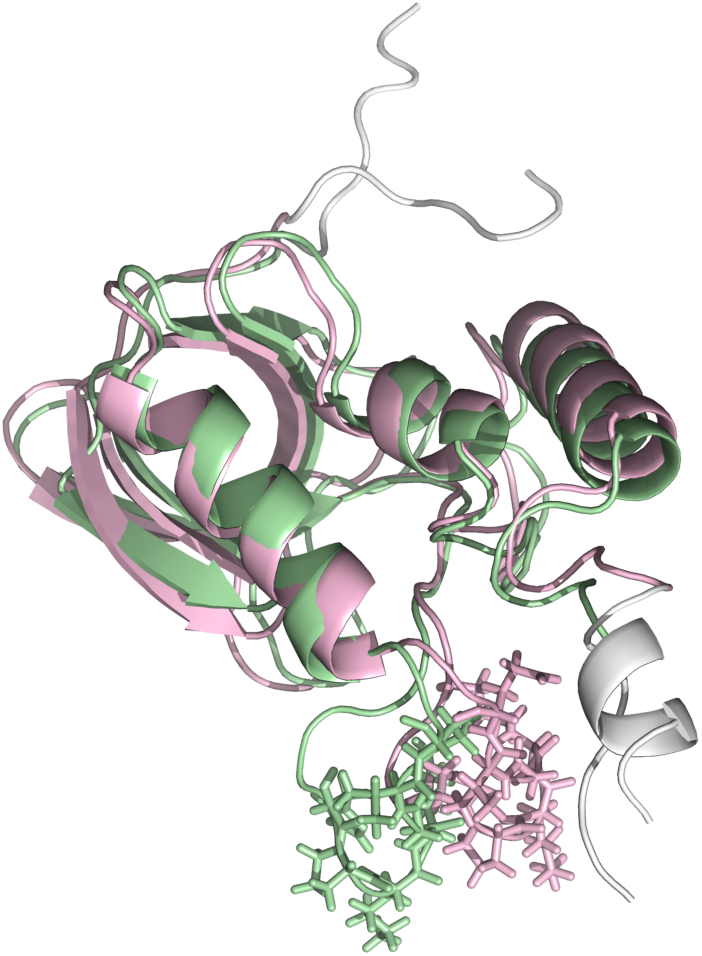
Snapshots of protein 31 at 110 ns (green) and 10001 ns (pink). The seven residues at the N- and C-terminus, ignored in the rotational fit, are colored white. The Euler angle ψ of the rotational fit decreases by 0.7 degrees when the tip of the flexible loop (residues 46-52, indicated using sticks) is removed. The image was rendered using Pymol (https://pymol.org/).

In Bragg analysis, a good molecular replacement solution yields a linear correlation with the Bragg data of about 0.80. The individual atom positions and B factors can then be refined to determine a crystal structure that is specific to the crystallography experiment. Because each Bragg reflection is determined by the entire crystal structure, local atomic details only become resolved once the entire structure is modeled with sufficient accuracy. Similarly, accurate models of diffuse data might only reveal the atomic details of molecular motions when the entire conformational ensemble is modeled with sufficient accuracy.

The present correlation of 0.68, although a significant advance, probably only reflects a global agreement of the model with the data, and not a validation of the details of the simulation. As for the Bragg data, once the correlation of models with the anisotropic diffuse data is sufficiently high, diffuse scattering figures of merit such as correlation coefficients or R factors might become more sensitive indicators of whether the MD motions are real. If this can be achieved, then crystallography and MD simulations will become a powerful tool for obtaining experimentally validated models of biomolecular mechanisms in crystalline proteins.

## Acknowledgments

This work was supported by the US Department of Energy, via the Exascale Computing Project. Many thanks to James S. Fraser, for suggesting the rigid-body motion analysis, and for leading the “Macromolecular movements by simulation and diffuse scatter” Laboratory Fees Research Project, which provided additional funding from the University of California. Thanks also to Peter B. Moore for reading the manuscript and suggesting the addition of the B factor comparison including the TLS refinement results. The simulations were performed using Institutional Computing machines at Los Alamos National Laboratory, supported by the US Department of Energy under Contract DE-AC52-06NA25396.

---

**Supplementary Figure S1.**
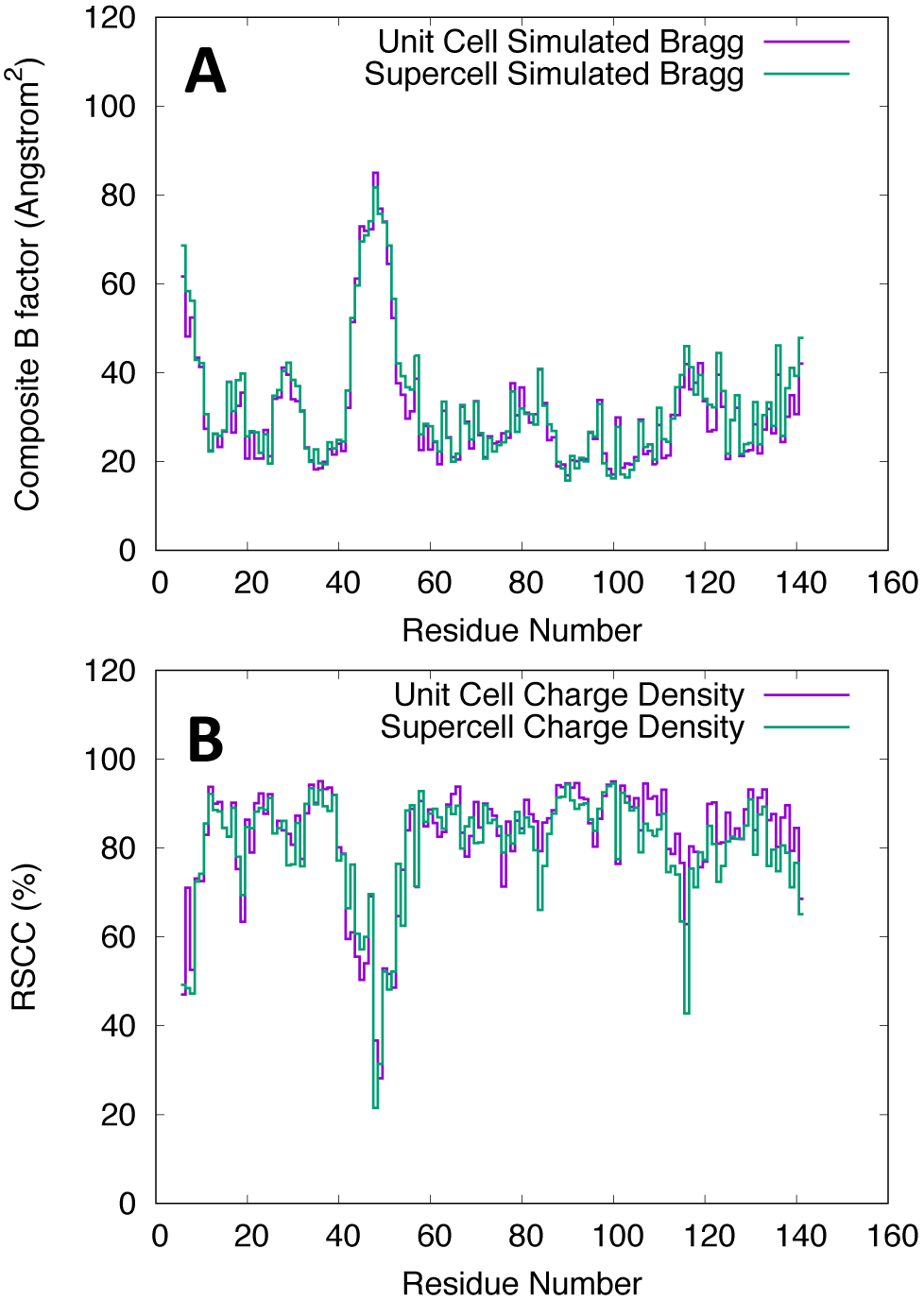
Comparison of Bragg analysis results for the unit cell vs. supercell simulations. (A) Residue-wise comparison between the B factors from the crystal structure (purple) and simulated average structures from the unit cell (green) and supercell (cyan) simulations. (B) Residue-wise RSCC computed between the crystal structure and the charge density from the supercell (green) and unit cell (cyan) simulations.

